# Cardioprotection by the mitochondrial unfolded protein response (UPR^mt^) is mediated by activating transcription factor 5 (ATF5)

**DOI:** 10.1101/344606

**Authors:** Yves T. Wang, Yunki Lim, Matthew N. McCall, Cole M. Haynes, Keith Nehrke, Paul S. Brookes

## Abstract

The mitochondrial unfolded protein response (UPR^mt^)^1^ is a cytoprotective signaling pathway triggered by mitochondrial dysfunction. Activation of the UPR^mt^ upregulates nuclear-encoded mitochondrial genes, including those for chaperones, proteases, and antioxidants, as well as glycolysis, to restore proteostasis and cell energetics. Activating transcription factor 5 (ATF5), a protein with both mitochondrial and nuclear targeting sequences, is proposed to mediate mammalian UPR^mt^ signaling. Since proteostasis and bioenergetics are important in the response of organs such as the heart to injury, we hypothesized that pharmacologic UPR^mt^ activation may be cardioprotective against ischemia-reperfusion (IR) injury and that such protection would require ATF5. Using a perfused heart IR injury model in wild-type and global *Atf5^−/−^* mice, we found that *in-vivo* administration of the UPR^mt^ inducers oligomycin or doxycycline 6 h prior to *ex-vivo* IR injury was cardioprotective. Such protection was absent in hearts from *Atf5*^−/−^ mice, and no protection was observed with acute *ex-vivo* cardiac administration of doxycycline. Loss of ATF5 also did not alter baseline IR injury (without UPR^mt^ induction). Cardiac gene expression analysis by RNA-Seq revealed mild induction of numerous genes in an ATF5-dependent manner, which may be important for cardioprotection. Analysis of hearts by qPCR showed that oligomycin at 6 h significantly induced genes encoding ATF5 and several known UPR^mt^-linked proteins. We conclude that ATF5 is required for cardioprotection induced by drugs that activate the UPR^mt^.

---

Tissue ischemia and hypoxia are important pathologic phenomena that underlie disease conditions such as myocardial infarction and stroke. Paradoxically, the reperfusion of ischemic/hypoxic tissues exacerbates injury, and ischemia-reperfusion (IR) injury causes significant morbidity and mortality worldwide (1). In the heart, mitochondria are key mediators of IR injury, and are thus a potential target for cardioprotective interventions (2, 3).

An important concept in many protective paradigms is hormesis, wherein small sub-lethal insults can trigger cell-signaling events that afford protection against subsequent larger injuries (4). “Mitohormesis” refers to such hormetic events at the mitochondrial level. A classic example is ischemic preconditioning (IPC), wherein small ischemic insults trigger mitochondrial generation of reactive oxygen species (ROS) that signal to induce protection against large-scale pathologic ROS generation during subsequent IR injury (3).

The mitochondrial unfolded protein response (UPR^mt^) is a mitohormetic stress response pathway triggered by proteotoxic stress in the organelle (5). Similar to other compartment-specific UPRs (e.g., the endoplasmic reticulum UPR), UPR^mt^ signaling upregulates chaperones to promote protein refolding and proteases to digest misfolded proteins (6). Furthermore, the UPR^mt^ also upregulates glycolysis (7) and down-regulates expression of mitochondrial respiratory chain subunits (8) to balance cell energetics and reduce burden on the mitochondrial translation and folding machinery.

While the UPR^mt^ was first discovered in mammals (5), significant insight to this pathway has emerged from work in *C. elegans*, including the discovery of its central transcription factor ATFS-1 (9). The ATFS-1 protein contains both nuclear and mitochondrial targeting sequences, and under normal conditions ATFS-1 is imported to mitochondria and degraded (7, 9). Under conditions of mitochondrial proteotoxicity, ATFS-1 import to the organelle declines, resulting in increased translocation to the nucleus and transcription of target genes (7, 8).

Comparatively little is known about the mammalian UPR^mt^, particularly in the heart. The transcription factor ATF5 was proposed as a mammalian ortholog of ATFS-1 (9), and indeed ATF5 can rescue UPR^mt^ signaling in *C. elegans* lacking ATFS-1 (10). Previously ATF5 was shown to have roles in cancer (11), cell cycle regulation (12), apoptosis (13), neural differentiation (14) and stress signaling (15). Importantly, to date there has been no *in-vivo* mammalian evidence that ATF5 is required for UPR^mt^ signaling.

Cardiac IR injury causes bioenergetic dysfunction, protein misfolding, and oxidative stress (16). The targets of UPR^mt^ signaling include glycolysis, chaperones, proteases, and antioxidants (6). Accordingly, we previously showed that UPR^mt^ induction was protective against anoxia-reoxygenation in *C. elegans* (17). Herein we hypothesized that induction of an ATF5-dependent UPR^mt^ may be cardioprotective. against IR injury in mammals.

The mitochondrial ATP synthase inhibitor oligomycin and the tetracycline antibiotic doxycycline are known to induce UPR^mt^ in mammalian cells (10, 18), likely via disruption of mitochondrial protein import and/or synthesis. Herein we tested the ability of these molecules to elicit cardioprotection in WT (*Atf5^+/+^*) and *Atf5^−/−^* mice (19), and we performed gene expression analysis to investigate ATF5-dependent signaling.

## Results

The complete original data set for the figures in this paper is deposited at https://www.figshare.com (DOI: 10.6084/m9.figshare.7323929).

### Baseline cardiac function and response to IR injury in Atf5^−/−^ mouse hearts

WT (*Atf5^+/+^*), *Atf5^+/−^*, and *Atf5^−/−^* mice (Fig. S1A/B) were used. Consistent with previous reports (19–21), major confounders in these studies were high neonatal mortality and reduced adult body weight of *Atf5^−/−^* mice (Fig. S1C/D). Thus, both sexes were used and controls were age- and sex-matched but not always littermates.

Langendorff-perfused *ex-vivo* hearts were subjected to global IR injury, with cardiac function monitored as rate × pressure product (RPP), and post-IR infarct size assayed by tetrazolium chloride (TTC) staining. *Atf5* genotype did not impact post-IR functional recovery or infarct size (Fig. 1A/B). Analysis with all genotypes aggregated showed no differences between males and females (Fig. 1C/D), and further stratification by both sex and genotype did not reveal any sex-specific genotype-dependent effects (Fig. 1 E/F). Analysis of baseline cardiac function stratified by both sex and genotype showed modest sex effects but no genotype effects (Fig. S2).

**Figure 1:**
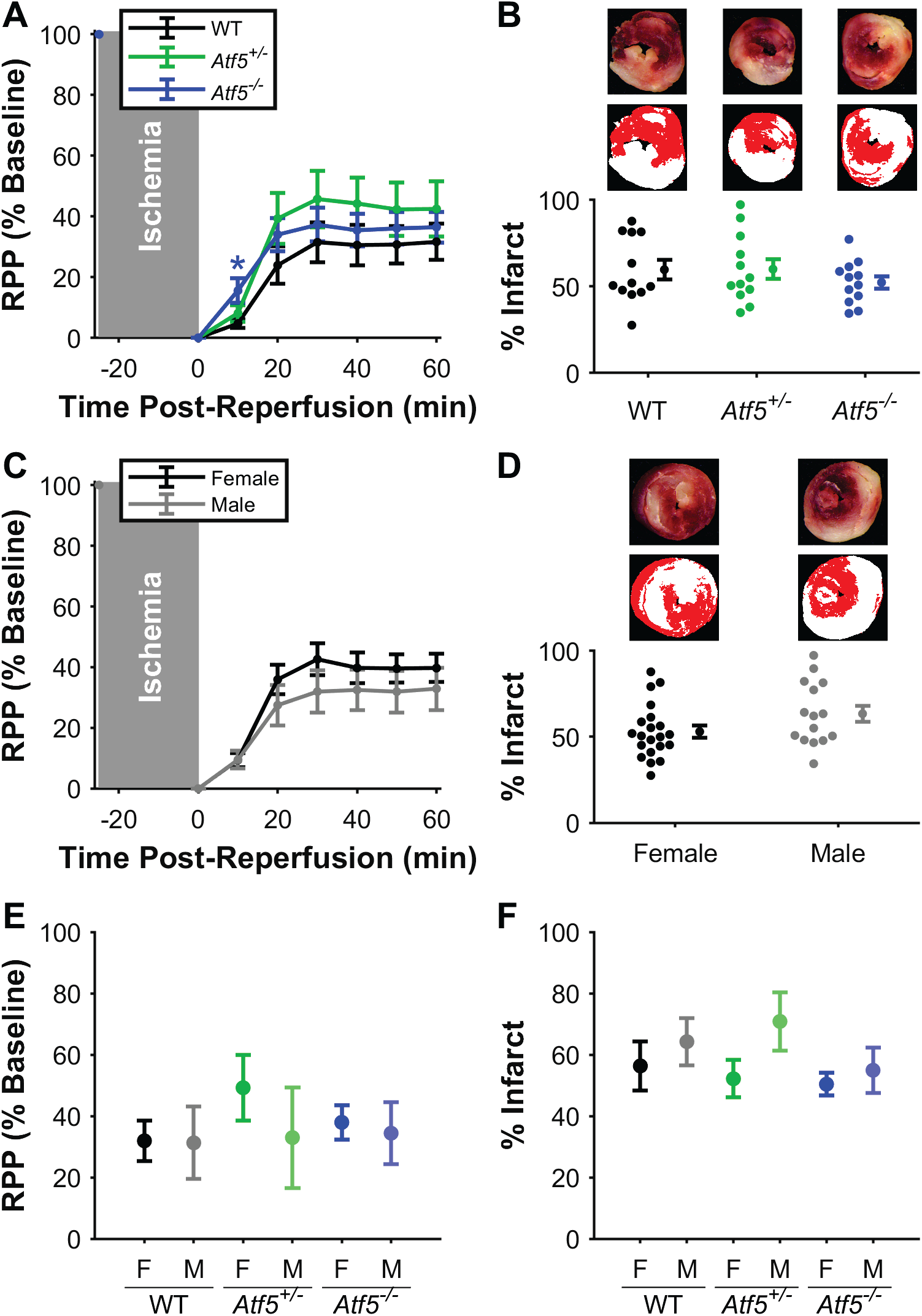
Absence of ATF5 does not affect vulnerability to IR injury. **(A)** Post-reperfusion functional recovery by genotype, measured by baseline-normalized rate × pressure product (RPP). Animals were of both sexes, n=12 per genotype (WT: black, *Atf5^+/−^*: green, *Atf5^−/−^*: blue). *p<0.05 vs. *Atf5^+/+^*. **(B)** Infarct sizes of the hearts in (A), measured by TTC staining (bottom), with representative stained slices (top) and segmented images (middle) with healthy tissue (red) and infarcted tissue (white). **(C)** Post-reperfusion functional recovery by sex. Hearts are the same as shown in (A) with mixed genotypes, n=21 for females (black) and n=15 for males (gray). **(D)** Infarct sizes of the hearts in (C). **(E/F)** Post-reperfusion functional recovery and infarct size for the same hearts stratified by both sex and genotype. Data are means±SEM.

### ATF5-dependent cardioprotection by oligomycin

Oligomycin induces the UPR^mt^ in mammalian cells (10), so we tested whether oligomycin administration prior to IR injury could elicit cardioprotection. Low-dose oligomycin was administered *in-vivo* (500 μg/kg) 6 h prior to *ex-vivo* IR injury. Hearts from WT oligomycin-mice exhibited significant improvement in post-IR functional recovery (oligomycin 52±4% vs. vehicle 24±4%, p=0.0018) and reduction in infarct size (oligomycin 39±3% vs. vehicle 64±9%, p=0.042) (Fig. 2A/B). This protective effect of oligomycin was absent in hearts from *Atf5^−/−^* mice (functional recovery: oligomycin 30±4% vs. vehicle 19±5%, p=0.18; infarct: oligomycin 58±6% vs. vehicle 57±8%, p=0.98) (Fig. 2C/D). Baseline cardiac function prior to IR was similar across groups (Fig. S3).

**Figure 2:**
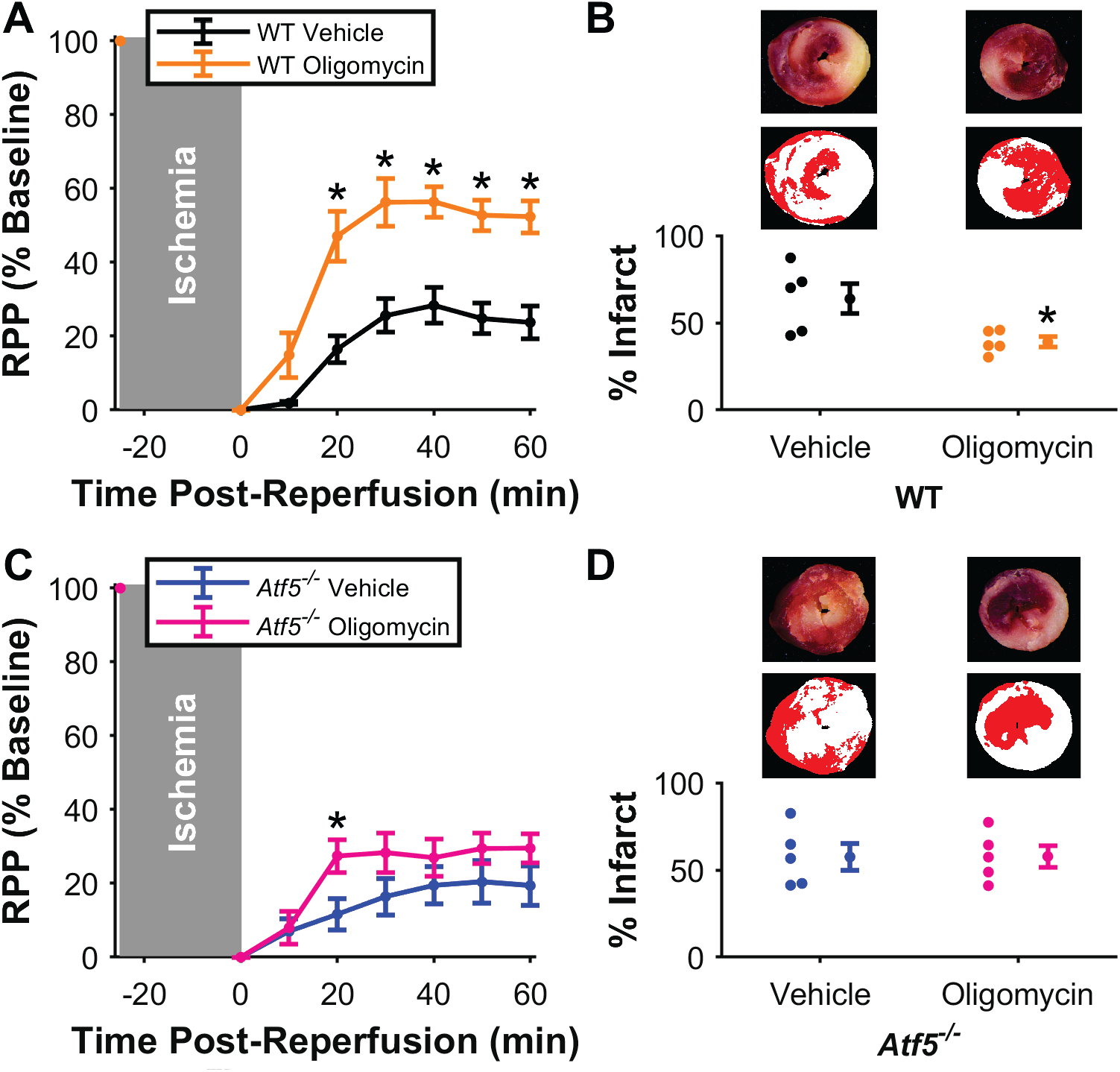
UPR^mt^ induction by oligomycin protects against IR injury and requires ATF5. **(A)** Post-reperfusion functional recovery, measured by baseline-normalized rate × pressure product (RPP) in WT mice given oligomycin (orange) or vehicle (black). n=5 per group. *p<0.05 compared to matching time point. **(B)** Infarct sizes of the hearts in (A), measured by TTC staining (bottom), with representative stained slices (top) and segmented images (middle) with healthy tissue (red) and infarcted tissue (white). *p<0.05 compared to vehicle control. **(C)** Post-reperfusion functional recovery of *Atf5^−/−^* hearts given oligomycin (pink) or vehicle (blue). n=5 per group. *p<0.05 compared to matching timepoint. **(D)** Infarct sizes of the hearts in (C). Data are means±SEM.

### ATF5-dependent cardioprotection by doxycycline

Doxycycline is known to elicit cardioprotection against IR injury, via a mechanism proposed to involve matrix metalloproteinases (MMPs) (22–25). Since doxycycline also disrupts mitochondrial ribosomal function (26–28) and activates the UPR^mt^ (18, 29), we hypothesized that it may elicit cardioprotection via the ATF5-dependent UPR^mt^.

Doxycycline was administered *in-vivo* (70 mg/kg) 6 h prior to *ex-vivo* IR injury. Hearts from WT doxycycline-treated mice exhibited significant improvement in post-IR functional recovery (doxycycline 60±8% vs. vehicle 23±4%, p=0.0052) and reduction in infarct size (doxycycline 37±3% vs. vehicle 69±4%, p=0.00046) (Fig. 3A/B). This protective effect of doxycycline was absent in hearts from *Atf5^−/−^* mice (functional recovery: doxycycline 42±7% vs. vehicle 34±7%, p=0.40; infarct: doxycycline 50±2% vs. vehicle 59±4%, p=0.11) (Fig. 3C/D). Baseline cardiac function prior to IR was similar across groups (Fig. S4).

**Figure 3:**
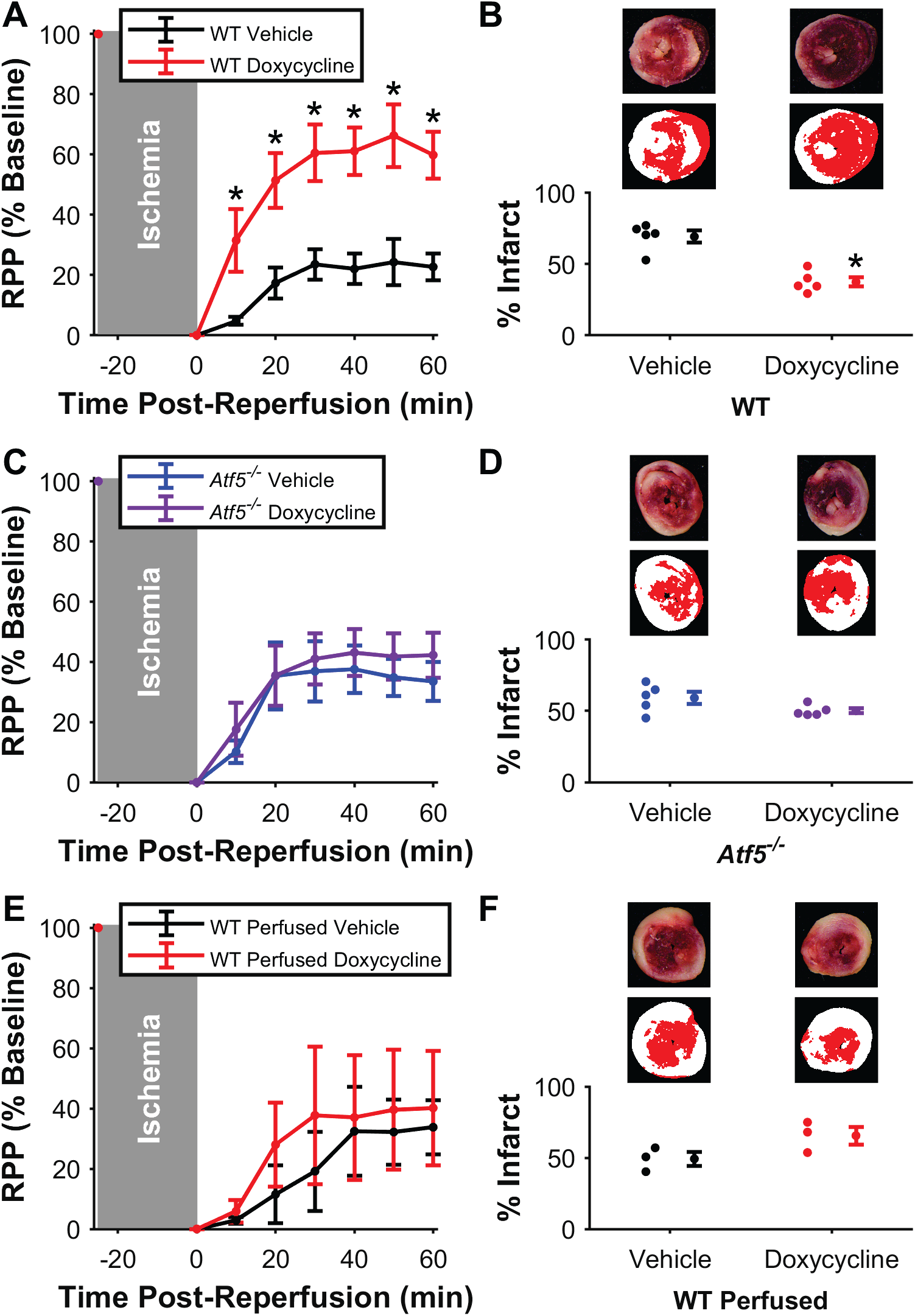
UPR^mt^ induction by doxycycline protects against IR injury and requires ATF5. **(A)** Post-reperfusion functional recovery, measured by baseline-normalized rate × pressure product (RPP) in WT mice given doxycycline (red) or vehicle (black). n=5 per group. *p<0.05 compared to matching time point. **(B)** Infarct sizes of the hearts in (A), measured by TTC staining (bottom), with representative stained slices (top) and segmented images (middle) with healthy tissue (red) and infarcted tissue (white). *p<0.05 compared to vehicle control. **(C)** Post-reperfusion functional recovery of *Atf5^−/−^* hearts given doxycycline (purple) or vehicle (blue). n=5 per group. **(D)** Infarct sizes of the hearts in (C). **(E)** Post-reperfusion functional recovery of WT hearts given acute doxycycline (red) or vehicle (black) *ex-vivo*. n=3 per group. **(F)** Infarct sizes of the hearts in (E). Data are means±SEM.

The ability of doxycycline to induce acute cardioprotection was also tested, by delivering the drug to *ex-vivo* perfused hearts prior to ischemia and during reperfusion. However, in contrast to previous reports (29), no significant protective effect was observed (functional recovery: doxycycline 40±19% vs. vehicle 34±9%, p=0.78; infarct: doxycycline 66±6% vs. vehicle 49±5%, p=0.11) (Fig. 3E/F).

### Genetic analysis of UPR^mt^ activation

To explore the cardioprotective signaling mechanism(s) of UPR^mt^ induction, we performed global gene expression analysis using RNA-Seq on cardiac mRNA from WT or *Atf5^−/−^* mice treated with vehicle, oligomycin, or doxycycline. Since both drugs are UPR^mt^ inducers, they were grouped for analysis to identify common genes induced by either drug in WT (adjusted p<0.2), with reduced or no induction in *Atf5*^−/−^ (adjusted p<0.4). Such analysis yielded 69 genes, indicating an ATF5-dependent response in this model. However, previously reported UPR^mt^-induced genes (30) were poorly represented in this gene set (Fig.4, Table S1). Furthermore, an unbiased pathway analysis did not yield useful insights to potential gene programs that may underlie the phenomenon of ATF5-dependent cardioprotection.

**Figure 4:**
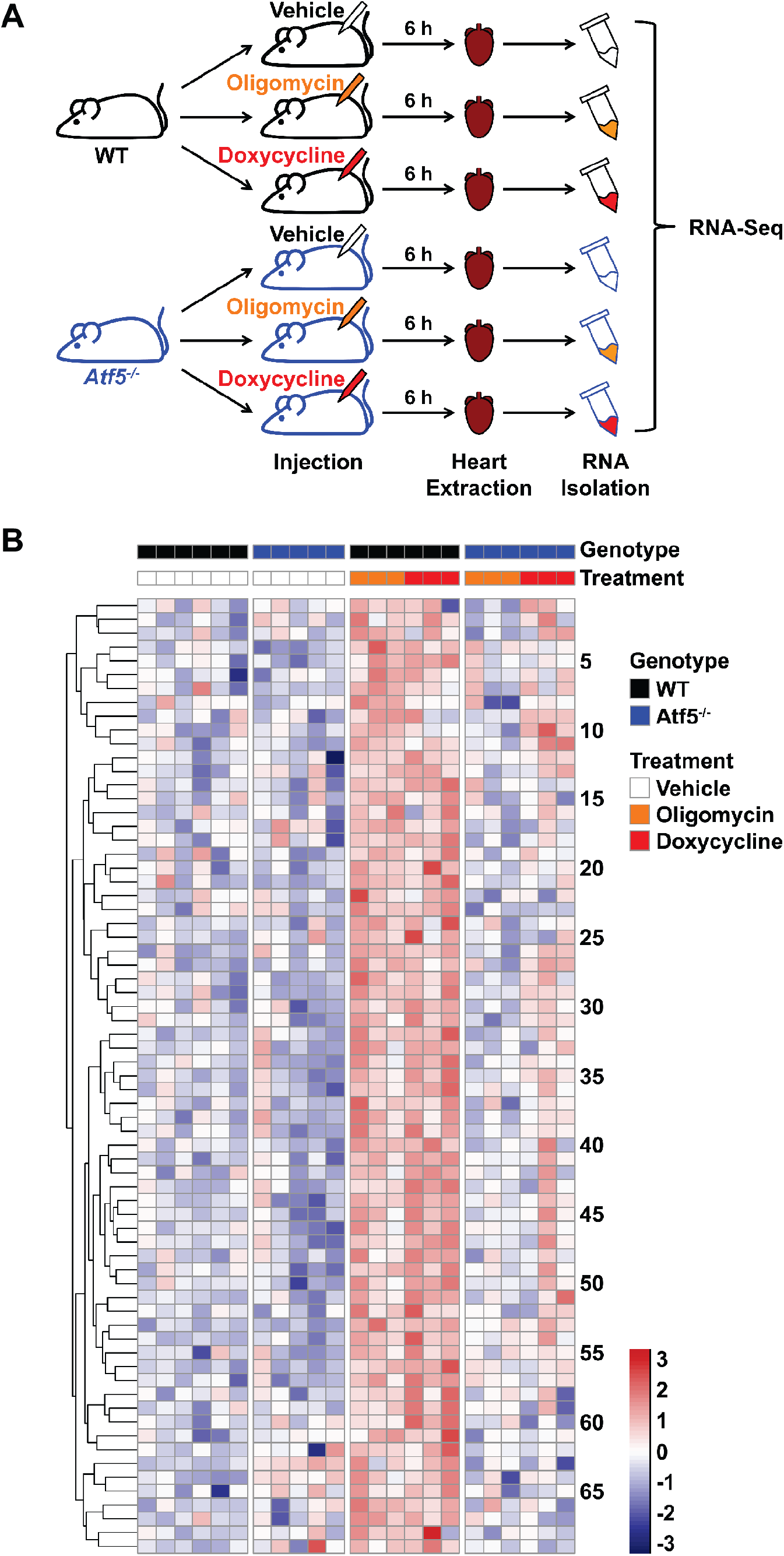
Gene expression analysis by RNAseq. **(A)** Schematic showing genotypes and treatment groups for RNA-Seq analysis on cardiac RNA extracts. **(B)** Heat map showing relative expression levels for the 69 genes identified as being significantly up-regulated by drug treatment (oligomycin or doxycycline) in WT mice and exhibiting a blunted response to drug treatment in *Atf5^−/−^*. Gene names are listed in Table S1. Full data set has been deposited in the Sequence Read Archive (SRA) under accession number SRP150238.

To probe the UPR^mt^ transcriptional response more directly, the expression of a set of known UPR^mt^ target genes was analyzed using RT-qPCR. Cardiac mRNA was isolated from WT mice 6 h after administration of oligomycin, and the change in expression due to oligomycin (vs. vehicle) was quantified relative to reference genes (Cq). As expected, the UPR^mt^-associated genes *Hspd1* (HSP60), *Clpp*, and *Lonp*, as well as *Atf5*, were significantly induced by oligomycin, providing evidence of a UPR^mt^-specific response in the heart. However, expression of *Hspa5* which encodes the chaperone BiP in the endoplasmic reticulum UPR was not induced, suggesting no global upregulation of chaperones or other UPR pathways.

## DISCUSSION

The key findings of this study are that pharmacologic induction of the UPR^mt^ is cardioprotective in IR injury and that ATF5 is necessary for this cardioprotection. Although a role for ATF5 in the UPR^mt^ has been proposed based on work in *C. elegans* and mammalian cell culture (10), this is the first *in-vivo* demonstration that ATF5 is a necessary component of the canonical UPR^mt^ in mammals. It is also the first study to demonstrate a role for the UPR^mt^ in cardioprotection by tetracycline antibiotics (22–25).

The mechanism(s) by which oligomycin or doxycycline activate the UPR^mt^ *in-vivo* are currently unclear. Since oligomycin inhibits mitochondrial ATP synthase, and protein synthesis generally ranks low in the hierarchy of ATP consuming processes in cells (31), it is possible that UPR^mt^ induction occurs via impairment of mitochondrial protein synthesis (32) or import (33), both of which are energy-intensive processes. Importantly, the dose of oligomycin used herein elicited no mortality and did not affect baseline cardiac function (Fig. S3). Doxycycline disrupts mitochondrial ribosomal function (26–28), and has been demonstrated to induce the UPR^mt^ in mammalian cells (10), presumably via producing an imbalance between nuclear and mitochondrial DNA encoded subunits of the respiratory chain (18).

A single bolus injection of oligomycin or doxycycline 6 h prior to IR elicited significant cardioprotection in WT mice, indicating that mitochondrial proteotoxic stress is a hormetic mechanism of cardioprotection, akin to other cardioprotective paradigms such as delayed preconditioning, wherein a primary stress induces a gene program that protects against subsequent similar stress (34). In the case of UPR^mt^, this gene program includes chaperones, proteases, antioxidants, and a shift away from mitochondrial oxidative metabolism toward glycolysis (6–8), all of which would be predicted to confer benefits during IR injury.

Cardioprotection conferred by pharmacologic UPR^mt^ induction was absent in *Atf5*^−/−^ mice. Importantly, there was no change in baseline IR injury with ATF5 genotype and no difference in baseline cardiac function between drug treatment groups, thus indicating the loss of cardioprotection in *Atf5*^−/−^ mice was not due to an increased sensitivity to IR injury or underlying effects of the UPR^mt^-inducing drugs on cardiac function.

Although RNA-Seq analysis led to the identification of several genes induced by either oligomycin or doxycycline in an ATF5-dependent manner, none of these were known UPR^mt^ targets. In contrast, qPCR identified a modest but significant increase in cardiac mRNA expression for several UPR^mt^-associated genes in oligomycin-treated mice. This discrepancy could be due to the low overall sensitivity of RNA-Seq relative to qPCR.

A potential confounder for the use of doxycycline to induce the UPR^mt^ in the context of cardioprotection is its reported ability to confer cardioprotection when administered post-IR injury via inhibiting MMPs (22–25). A human clinical trial was based on such findings (NCT00469261), and the proposed mechanisms for MMP inhibition by doxycycline include divalent metal chelation (35) and redox activity (36). However, our data indicate that doxycycline-induced cardioprotection requires ATF5. To the best of our knowledge there is no connection between ATF5 and MMPs, and no MMP genes emerged from our RNA-Seq analysis. Furthermore, the structurally unrelated UPR^mt^-inducer oligomycin also yielded ATF5-dependent cardioprotection, suggesting the UPR_mt_/ATF5 axis is a standalone mechanism of doxycycline inducible cardioprotection, independent from MMPs.

Although herein we chose a 6 h delay between drug administration and IR injury, to permit induction of the UPR^mt^ gene program, a recent report claimed acute cardioprotection by tetracycline antibiotics delivered 30 min prior to ischemia (29). Our inability to reproduce this phenomenon may have been due to an observed low solubility of doxycycline in Krebs-Henseleit perfusion buffer, such that the microfiltration necessary for successful mouse cardiac perfusion (37) may have limited doxycycline bioavailability in our experiments. In addition, a 30 min pre-treatment is likely too short for induction of a genetic stress response such as the UPR^mt^ (38). Thus, acute tetracycline cardioprotection is likely attributable to MMP-inhibition, as previously shown (22).

The current results are the first to cement a role for ATF5 as the central mediator of the canonical UPR^mt^ in mammals at the whole animal level. While the UPR^mt^ has been extensively characterized in *C. elegans*, it remains unclear whether the mammalian UPR^mt^ retains all features observed in the model organism. For example, the nematode UPR^mt^ involves several proteins with potential but unproven orthologs in mammals, including DVE-1 (SATB2) and UBL-5 (UBL5) (39). Another key feature of the UPR^mt^ uncovered in *C. elegans* is its potential for cell non-autonomous signaling, whereby UPR^mt^ induction in one cell or organ can trigger protection against stress in another (40). In this regard, we observed mild induction of the UPR^mt^ in hearts of oligomycin-treated animals, but it should be noted that we utilized a global *Atf5*^−/−^ mouse. As such, we cannot exclude the possibility that pharmacologic ATF5-dependent UPR^mt^ induction *in-vivo* may have occurred in a tissue other than the heart, with a blood-borne or other signal eliciting remote cardioprotection via ATF5-independent mechanisms. Further studies utilizing tissue-specific *Atf5^−/−^* animals may therefore prove insightful.

## Experimental procedures

### Reagents, animals

All reagents were from Sigma (St. Louis, MO) unless otherwise stated. All animal experiments were approved by the Committee on Animal Resources of the University of Rochester (Protocol #UCAR-2014-036) in compliance with the NIH *Guide for the Care and Use of Laboratory Animals*: 8^th^ edition (revised 2011). Founder *Atf5^−/+^* mice were provided by Stavros Lomvardas on a C57BL/6(J/N) background (19) and backcrossed to C57BL/6J (000664) from JAX (Bar Harbor, ME) at least three generations. Mice were housed with a 12-hour light/dark cycle, with *ad libitum* food and water. Experimental mice were 12-20 week adults of both sexes (n=131). WT (*Atf5^+/+^*) controls were age- and sex-matched, as the high neonatal mortality of *Atf5^−/−^* mice made using littermates infeasible. PCR genotyping was performed at 21 days by analyzing tail clip DNA with a mix of 3 primers, with expected amplicon sizes of 1006bp for the WT allele and 439bp for the knockout allele:

WT: 5’-TCTGATTGGATGACTGAGCGG-3’
KO: 5’-GCAGCCTCTGTTCCACATACACTTC-3’
Common: 5’-TCACTTGTGTTCCAAGTCCCC-3’

### Perfused heart IR injury model

IR injury was assessed using an *ex-vivo* Langendorff perfused heart model as previously described (41). Following intraperitoneal administration of anesthetic (tribromoethanol, 200 mg/kg) and heparin (0.2 mL), each mouse was placed supine on a 37°C warming pad, and the aorta cannulated with a blunt 37 G needle followed by immediate transfer to the perfusion rig. Perfusion was at 4 mL/min with 37°C, 0.22μm filtered, Krebs-Henseleit buffer comprising: 118 mM NaCl, 4.7 mM KCl, 1.2 mM MgSO_4_, 25 mM NaHCO_3_, 1.2 mM K_2_HPO_4_, gassed with 95% O_2_/5% CO_2_. Additionally 5 mM glucose, 100 μM palmitate-BSA, 200 μM pyruvate, and 1.2 mM lactate were present as metabolic substrates. Left ventricular (LV) pressure was measured by a transducer-linked water-filled LV balloon, with digital recording (DATAQ, Akron, OH). LV pressure traces were analyzed using a custom MATLAB program (MathWorks, Natick, MA), to calculate developed pressure (dP), heart rate (HR), rate × pressure product (RPP), and maximal rate of rise (+dP/dt_max_). Hearts were equilibrated for 25 min, then subjected to 25 min global no-flow ischemia followed by 60 min reperfusion. Post-IR hearts were transverse sliced (2 mm thick), stained with 1% tetrazolium chloride (TTC), fixed in formalin for 24 h, and slices imaged. Images were digitally segmented (healthy red tissue vs. infarcted white tissue) and pixel counts quantified using a custom MATLAB program.

### Drug Treatments

For *in-vivo* experiments, oligomycin (Sigma O4876, 500 μkg) or doxycycline (Sigma D9891, 70 mg/kg) were administered via intraperitoneal injection in a vehicle comprising of sterile normal saline with 0.1% DMSO (injection volume 12.5 μL/g body weight). Controls received vehicle alone. Mice were returned to cages for 6 h prior to experiments. For *ex-vivo* experiments, doxycycline (20 μM) was dissolved in perfusion buffer containing 0.25% DMSO, followed by 0.2μm filtration. Doxycycline-containing buffer was perfused throughout pre-ischemic equilibration and reperfusion.

### RNASeq and qPCR

Following anesthesia (tribromoethanol, 200 mg/kg) hearts were excised, flushed with ice-cold saline, homogenized in TRIzol reagent (Invitrogen, Carlsbad, CA), snap-frozen (liquid N_2_) and stored at −80°C overnight. RNA was processed from TRIzol according to the manufacturer’s instructions and further purified using a RNeasy Minikit (Qiagen, Hilden, Germany) coupled with on-column DNase treatment.

For RNA-Seq, library preparation and analysis were performed by the Functional Genomics Core at the University of Rochester (oligomycin experiments) or the Integrated Genomics Operation at Memorial Sloan Kettering Cancer Center (doxycycline experiments). Total RNA concentration was determined spectrophotometrically (NanoDrop 1000, Wilmington, DE) and RNA quality assessed with the Agilent Bioanalyzer (Agilent, Santa Clara, CA). Library construction and next-generation sequencing was performed from 200 ng RNA with the TruSeq Stranded mRNA Sample Preparation Kit and a HiSeq 2500 v4 platform with cBot according to manufacturer’s protocols (Illumina, San Diego, CA). Raw reads were demultiplexed using bcl2fastq.pl v1.8.4 conversion software (Illumina). Quality filtering and adapter removal were performed using Trimmomatic version 0.32 (42), then processed/cleaned reads were mapped to the mouse genome with STAR 2.4.2a (43).

For qPCR, cDNA was generated using an iScript cDNA Synthesis Kit (Bio-Rad, Hercules, CA) and used at 100 μg/well as a PCR template for target detection with iTaq Universal SYBR Green Supermix (Bio-Rad) on a CFX Connect Real-Time PCR Detection System (Bio-Rad). Quantification cycle (Cq) of each RT-qPCR targets was normalized to that of the reference genes actin (*Actb*) or *Hprt* (Cq). Primers and expected amplicon sizes are in Table S2. Data are presented in treated vs. untreated (vehicle) conditions as relative Cq (i.e., Cq).

### Statistics

Comparisons between 2 groups were performed using two-tailed, unpaired Student’s *t*-tests. Comparisons versus zero (Fig. 5) were performed by two-tailed, one-sample Student’s *t*- tests. Univariate comparisons between more than 2 groups (Fig. 1A/B) were performed by one-way ANOVA. Multivariate comparisons between groups (Fig. 1E/F, S2, S3, S4) were performed by two-way ANOVA. In all cases, differences were considered significant for p<0.05. Data are reported as means±SEM.

**Figure 5:**
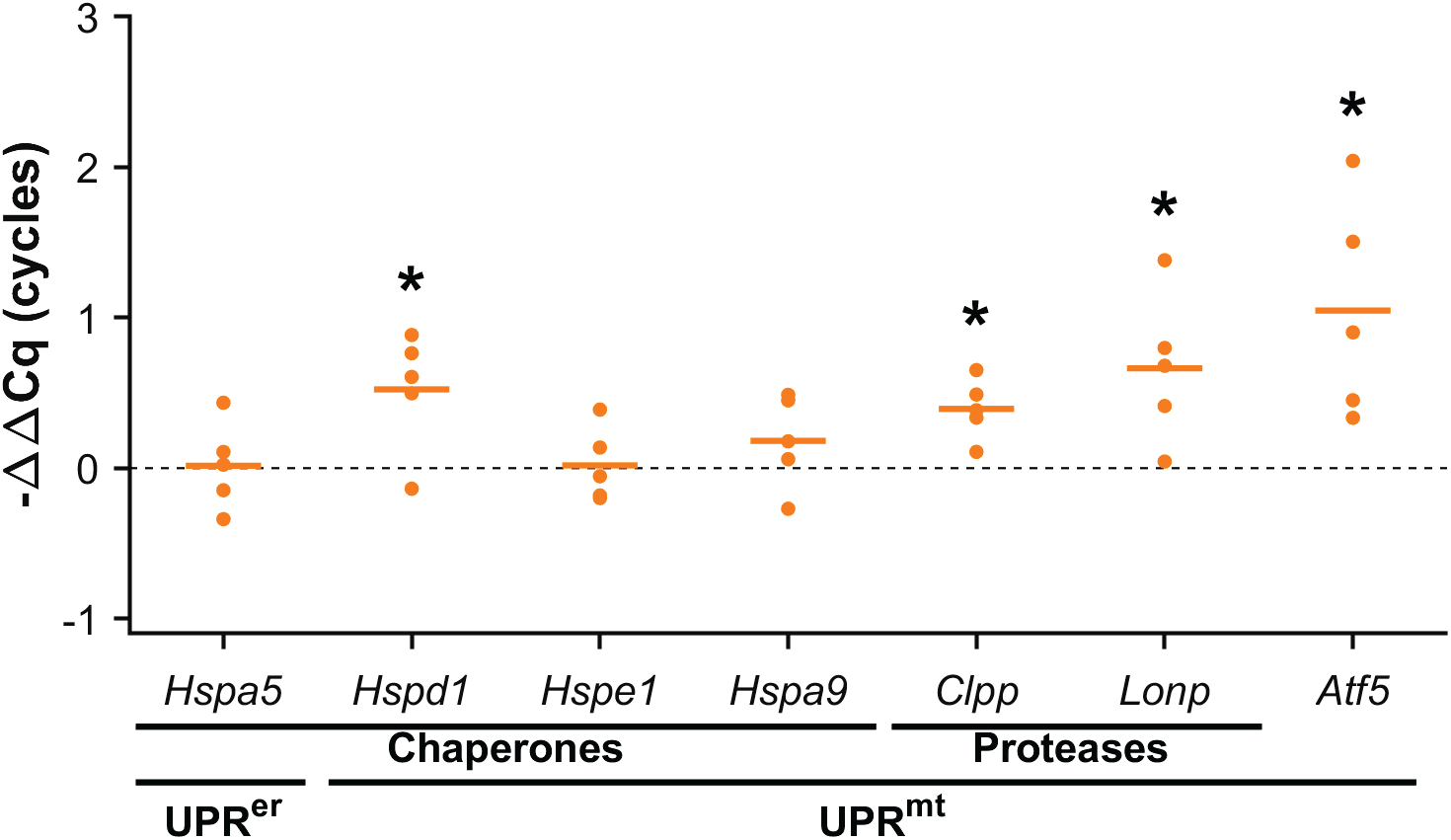
qPCR analysis of UPR^mt^ target genes induced by oligomycin. RT-qPCR was used to monitor relative expression of selected genes in hearts 6 h after treatment with oligomycin (− Cq, normalized to *Hprt1* reference gene and compared to vehicle-treated hearts). *Hspa5* (BiP) is an endoplasmic reticulum UPR chaperone. *Hspd1* (HSP60), *Hspe1* (HSP10), and *Hspa9* (mtHSP70) are UPR^mt^ chaperones. *Clpp* and *Lonp* are UPR^mt^ proteases. *Atf5* is the putative master transcriptional regulator of the UPR^mt^. Horizontal lines represent means. *p<0.05 compared to 0, n=5 per group.

RNA-Seq gene read counts were processed using the open source R/Bioconductor software suite (R version 3.4.4; Bioconductor version 3.7; *limma* version 3.34.9). One sample from an *Atf5^−/−^* mouse which received vehicle was removed prior to analysis because it contained reads that mapped to the *Atf5* gene indicating that it may not have been a true knockout. Genes with fewer than 50 reads in all samples were filtered prior to analysis, yielding 14,061 genes used in all subsequent analyses. The voom method (44) was used to estimate the mean-variance relationship of the logged gene counts and calculate a precision weight for each observation. These weights were then used with *limma* empirical Bayes analysis methodology (45) to assess differential expression. The following linear model was the basis of all statistical analyses:

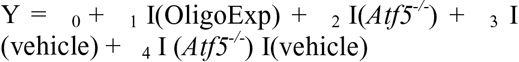

OligoExp includes the data from oligomycin-treated animals and their vehicle-treated controls. Gene set testing was performed using ROAST (46), a method that appropriately handles correlation between genes. RNA-Seq data have been deposited in the Sequence Read Archive (SRA) under accession number SRP150238.

## Acknowledgements

We thank Stavros Lomvardas (Columbia University, NY) for providing *Atf5^+/−^* founders. This work was supported by NIH R01-HL127891 and R01-HL071158.

## Footnotes

1 The abbreviations used are: UPR^mt^, mitochondrial unfolded protein response; ATF5, activating transcription factor 5, IR, ischemia-reperfusion; IPC, ischemic preconditioning; ROS, reactive oxygen species; RPP, rate × pressure product; TTC, tetrazolium chloride; MMP, matrix metalloproteinase; LV, left ventricular

